# Probabilistic ancestry maps: a method to assess and visualize population substructures in genetics

**DOI:** 10.1101/362343

**Authors:** Héléna A. Gaspar, Gerome Breen

## Abstract

Principal component analysis (PCA) is a standard method to correct for population stratification in ancestry-specific genome-wide association studies (GWASs) and is used to cluster individuals by ancestry. Using the 1000 genomes project data, we examine how non-linear dimensionality reduction methods such as t-distributed stochastic neighbor embedding (t-SNE) or generative topographic mapping (GTM) can be used to provide improved ancestry maps by accounting for a higher percentage of explained variance in ancestry, and how they can help to estimate the number of principal components necessary to account for population stratification. GTM also generates posterior probabilities of class membership which can be used to assess the probability of an individual to belong to a given population - as opposed to t-SNE, GTM can be used for both clustering and classification. This paper is a first application of GTM for ancestry classification models. Our maps and software are available online.

**Author summary:** With this paper, we seek to encourage researchers working in genetics to use other methods than PCA to visualize ancestry and identify substructures in populations. We propose to use methods which do not only allow visualization of ancestry, but also the estimation of probabilities of belonging to different ancestry groups.

## Introduction

As of 2018, most genome-wide association studies (GWASs) have used populations of European ancestry. However, larger sample sizes are now available and both societal need and funders are mandating more studies focused on other populations. Visualizing and accurately defining complex population structure is therefore of paramount importance. In this paper, we have three aims: to find a better way to visualize population substructures, to define a new procedure to estimate the optimal number of principal components accounting for population stratification, and to obtain an ancestry classification algorithm which can also estimate probabilities to belong to different ancestry groups. This paper focuses on global (genome-wide) ancestry rather than local ancestry defined within chromosome segments.

Principal component analysis (PCA) is widely used to investigate population structure in genetics [1], and to account for population stratification in GWASs (cf. EIGENSTRAT software [2]). However, the 2 or 3 principal components used to build a PCA plot generally account for a small percentage of variance explained and lead to a simplified visualization of population substructures, focused on major continental ancestry, with only partial sensitivity for the identification of admixed individuals or more complex ancestry. Model-based methods such as STRUCTURE [3] and ADMIXTURE [4] provide maximum likelihood estimations of ancestry based on ancestry proportions and allele frequencies but do not provide a visualization of population clusters as PCA and other multivariate analysis methods do.

A PCA ancestry map is constructed from a genotype matrix **G** of dimension *N × D*, where the *N* instances are individuals and the *D* features correspond to genetic variants - typically SNPs which are pruned to remove SNPs in high linkage disequilibrium with each other so that the identified principal components do not reflect local haplotype structure, but instead reflect genome-wide ancestry. For example, *G_nd_* could be the minor allele count for SNP *d* in individual *n*. For visualization purposes, PCA is used to map **G** to a more interpretable latent or hidden space of 2 or 3 dimensions: **G → X**, where X has dimension *N* × 2 or *N* × 3. The new variables - typically two for a PCA plot - are the first principal components, which account for the highest percentage of the overall variance. However, the total percentage of variance explained by such a small number of principal components can be low for high-dimensional genotype matrices.

More complex visualization methods such as t-distributed stochastic neighbor embedding (t-SNE) [5] or generative topographic mapping (GTM) [6], which are manifold-based and non-linear dimensionality reduction algorithms, are able to capture more information by embedding a *D*-dimensional space in a low-dimensional latent space, where *D* can be any number of features. Instead of two or three principal components, any number of principal components can be used with these methods. To assess the percentage of variance to account for population substructures, we propose to execute two mappings, first carrying out PCA to select principal components and then using t-SNE or GTM: **G → X’ → X**, where **X’** is the matrix of *F* principal components (*F* > 2), and **X** is the final t-SNE or GTM projection in a 2-dimensional space. The performance of ancestry classification models built with **X** or the visual assessment of clusters in **X** could then provide a way to estimate the number of principal components to account for population stratification.

Both t-SNE and GTM are used for clustering tasks. However, new instances cannot be projected onto a t-SNE map without training the map once again. GTM, on the other hand, not only allows for the projection of new data points, but comes with a probabilistic framework to build a comprehensive classification model and assign probabilities of class membership. GTM has been used in cheminformatics to classify compounds based on a simple map [7], to compare chemical libraries, and to assess their similarities [8]. GTM could easily be transposed to genetics and used to predict ancestry and degree of admixture in an individual or a group.

In this paper, 1000 Genomes Project Phase III [9] data is used to build the genotype matrix **G**. The 1000 Genomes Project has gathered genotypes from 26 different populations corresponding to 5 superpopulations: Africans (AFR), Admixed Americans (AMR), East Asians (EAS), Europeans (EUR) and South Asians (SAS). Ancestry maps are investigated to cluster and visualize superpopulations and populations using PCA, t-SNE, and GTM. t-SNE and GTM maps accounting for 3 to 1000 principal components are compared to a simple PCA plot. We also compare GTM ancestry classification models to two different algorithms: k-nearest neighbors (*k*-NN) models based on the 2D PCA plot, and linear Support Vector Machine (SVM), a classical machine learning algorithm [10]. We also demonstrate how to assess probabilities of ancestry membership in individuals and populations using GTM.

## Materials and methods

### Data and quality control

Genotypes of 2504 people in the 1000 Genomes Project Phase III were downloaded from ftp://ftp.1000genomes.ebi.ac.uk/vol1/ftp/release/20130502 [9]. Variants were removed based on a Hardy-Weinberg equilibrium (HWE) exact test p-value filter (< 0.001) and missing call rate filter (*>* 0.02). The HWE test measures whether the ratio between homozygous and heterozygous genotypes differs significantly from prediction under HWE assumptions. SNPs from the major histocompatibility complex (MHC) on chromosome 6 and in the chromosome 8 inversion region were excluded. The remaining SNPs were pruned twice using plink 1.9 [11, 12] with windows of 1000 variants and step size 10, pair-wise squared correlation threshold = 0.02, and minor allele frequency *>* 0.05. The pruning operation deals with *linkage desequilibrium* or non-random association of alleles at different loci: it reduces the number of SNPs, keeps SNPs in linkage equilibrium, and thereby reduces data dimensionality. A training set was built by removing the following populations: Americans of African ancestry in South West USA (code = ASW); African Caribbeans in Barbados (ACB); Mexican ancestry from Los Angeles USA (MXL); Gujarati Indian from Houston, Texas (GIH); Sri Lankan Tamil from the UK (STU); and Indian Telugu from the UK (ITU). We used these populations as an external test set to predict the degree of admixture in individuals and populations. For the classification models, we also merged British in England and Scotland (GBR) and Utah Residents with Northern and Western European Ancestry (CEU) to obtain a single category for Northern and Western European Ancestry.

### Visualization of ancestry clusters using PCA, t-SNE and GTM

t-SNE [5] translates similarities between points into probabilities; Gaussian joint probabilities in the original input space and Student’s t-distributions in the latent space. The Kullback-Leibler divergence between data distributions in the input and latent space is minimized with gradient descent. t-SNE has several parameters to optimize: the learning rate for gradient descent, the perplexity of distributions in the initial space, and the early exaggeration. However, the algorithm is only moderately sensitive to these two last parameters. In this paper, we used the scikit-learn implementation for t-SNE [13], with default learning rate = 200, perplexity = 30, and early exaggeration = 12. The main disadvantage of t-SNE is its lack of a framework to project new points onto a pre-trained map - a feature available in PCA and GTM.

The core principle of GTM [6] is to fit a manifold into the high-dimensional initial space. The points **y**_*k*_ on the manifold **Y** in the initial space are the centers of normal probability distributions of **g**, which here are individuals described by the genotype matrix **G**:

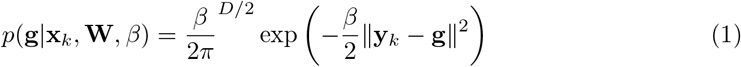

where *β* is the common inverse variance of these distributions and **W** is the parameters matrix of the mapping function **y**(**x**; **W**) which maps nodes **x**_*k*_ in the latent space to **y**_*k*_: **y**(**x**_*k*_; **W**) = **W***ϕ*(**x**_*k*_), where *ϕ*(**x**_*k*_) is a set of radial basis functions. **W** and *β* are optimized with an expectation-maximization (EM) algorithm maximizing the overall data likelihood. The responsibility or posterior probability that the individual **g**_*n*_ in the original genotype space is generated from the *k*th node in the latent space is computed using Bayes theorem:

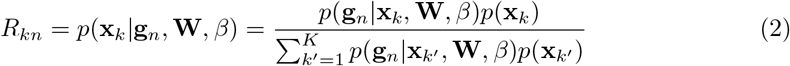

These responsibilities are used to compute the mean position of an individual on the map **x**(**g**_*n*_), by averaging over all nodes on the map:

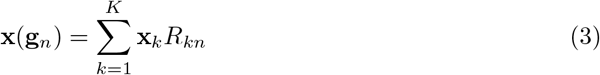

We used the python package ugtm v1.1.4 [14] for generative topographic mapping (https://github.com/hagax8/ugtm). The github wiki provides a tutorial for ancestry classification (https://github.com/hagax8/ugtm/wiki/2.-ugtm-for-ancestry-clustering). GTM has several hyperparameters to tune, which might have a high impact on the shape of the map: the number of radial basis functions, a width factor for these functions, map grid size, and a regularization parameter.

### Ancestry classification models

PCA does not provide a comprehensive framework to build a probabilistic classification model. However, a simple classification model associated with the 2-dimensional plot can be built using the *k*-NN approach in three steps: (1) a PCA plot is constructed from a training set, (2) a test set is projected on the plot, and (3) each test individual is assigned the predominant ancestry amongst its *k* nearest neighbors in the training set. We did not construct *k*-NN models for t-SNE since it is not straightforward to project new points onto a t-SNE map. On the other hand, GTM provides a probabilistic framework which can be used to build classification models and generate class membership probabilities [7]. GTM responsibilities can be seen as feature vectors: they encode individuals depending on their position on the map, which is discretized into a finite number of nodes (positions). They can be used to estimate the probability of a specific ancestry given the position on map, using Bayes’ theorem

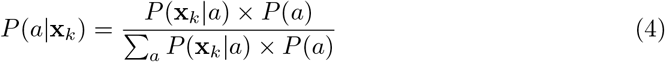

where *P*(**x***_k_*|*a*) is computed as follows:

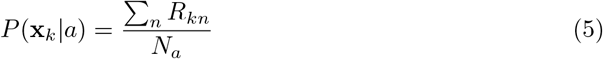

where *R_kn_* is the responsibility of node **x**_*k*_ for an individual belonging to population *a*, which counts *N_a_* individuals. It is then possible to predict the ancestry profile *P* (*a|***g**_*i*_) of a new individual with associated responsibilities {*R_ki_*}

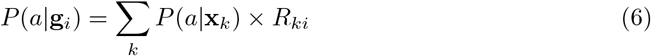

GTM nodes **x**_*k*_ can be represented as points colored by most probable ancestry *a*_max_ using *P*(*a*_max_ | **x**_*k*_). We compared performances of visual classifications (PCA and GTM) with linear support vector machine classification (SVM), a classical machine learning algorithm. Linear SVM is only dependent on *C*, the penalty hyperparameter. Increasing *C* increases the variance of the model and decreases its bias. In this application, predicted for 5 partitions of the data, which are concatenated to obtain predicted values for the entire dataset. From these, F1 scores are computed for each class *a* and repetition *j*. The per-class performance measure is computed across the 10 repetitions:

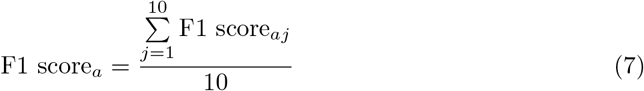

The overall model performance measure is a weighted average across per-class F1 scores, with weights equal to the number of individuals in the class:

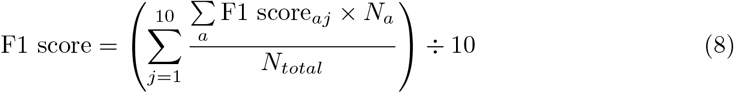

This procedure is performed for each parameter combination and for each algorithm (PCA, GTM, SVM). The best model for each algorithm is defined as having the largest overall F1 score. For PCA, we vary *k* (the number of neighbours) from 1 to 10. For GTM, we set the map grid size (number of nodes) = 16*16, the number of RBFs = 4*4, regularization = 0.1 and rbf width factor = 0.3. For linear SVM, the penalty parameter is set to *C* = 2^*r*^ where *r* runs from −5 to 10.

### Posterior probabilities of ancestry membership for whole populations

All our models are trained with only twenty 1000 Genomes Project populations. Six populations are used as an external test set (cf. foregoing section Data and quality control). Posterior probabilities of ancestry membership are estimated for all individuals in these test populations (Eq 6) based on observed superpopulation distributions (Eq 5). We also generate probabilities of belonging to a superpopulation for each population as a whole, by replacing individual responsibilities {*R_ki_*} in equation 6 by an overall population responsibility {*R_kp_*}

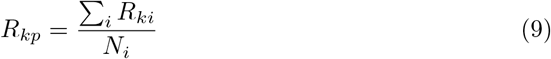

It should be noted that these responsibilities {*R_kp_*} correspond to the averaged distribution of the population on the map, and can be used to compare populations and estimate their diversity.

## Results and Discussion

### Classification of 5 superpopulations

PCA clusters and predicts the 5 superpopulations efficiently (F1 score = 0.98, cf. Table 1): Europeans, Africans, South Asians, East Asians, and Admixed Americans. However, SVM and GTM models with 3 or 10 principal components have higher recall for Admixed Americans and higher precision for South Asians. Optimal performances can be achieved by including a third principal component.

**Table 1.** 10 times repeated 5-fold cross-validated F1 score in five 1000 Genomes Project superpopulations using SVM, PCA or GTM.

| Ancestry | 1000G code | PCA 8-NN | SVM 10 PCs | GTM 3 PCs | GTM 10 PCs |
| --- | --- | --- | --- | --- | --- |
| Africans | AFR | 1.00 ± 0.00 | 1.00 ± 0.00 | 1.00 ± 0.00 | 1.00 ± 0.00 |
| Admixed Americans | AMR | 0.93 ± 0.00 | 1.00 ± 0.00 | 1.00 ± 0.00 | 1.00 ± 0.00 |
| East Asians | EAS | 1.00 ± 0.00 | 1.00 ± 0.00 | 1.00 ± 0.00 | 1.00 ± 0.00 |
| Europeans | EUR | 0.99 ± 0.00 | 1.00 ± 0.00 | 1.00 ± 0.00 | 1.00 ± 0.00 |
| South Asians | SAS | 0.93 ± 0.01 | 1.00 ± 0.00 | 1.00 ± 0.00 | 1.00 ± 0.00 |
| Overall F1 score |  | 0.98 ± 0.00 | 1.00 ± 0.00 | 1.00 ± 0.00 | 1.00 ± 0.00 |

SVM10 = support vector machine classification model using 10 principal components, PCA = k-nearest neighbours model based on 2D PCA map (k = 7), GTM{3,10,100} = bayesian classification model based on generative topographic mapping using 3, 10 or 100 principal components.

**Fig 1.**
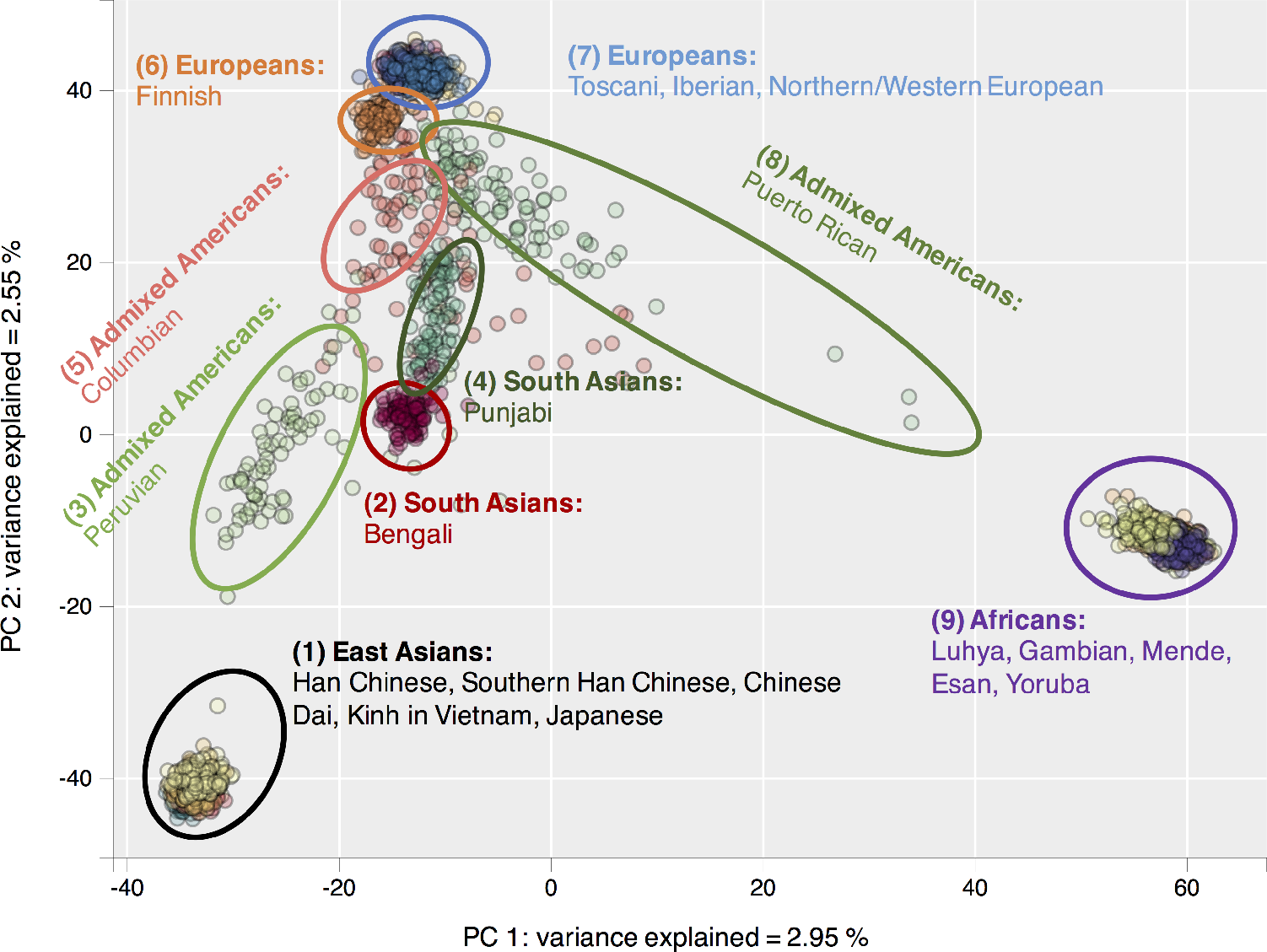
PCA clustering. PCA plot of 20 populations from 1000 Genomes Project, built using 2 first principal components.

### Maps for 20 populations using PCA, t-SNE and GTM

From Fig 2 and Fig 3, it can be seen that t-SNE and GTM recognize the same clusters. However, t-SNE provides clearer cluster visualization, whereas GTM suffers from a packing effect, which results in data points between packed together on a map. t-SNE remedies this situation with Student’s t-distributions in the latent space, which allow small distances between data points in the original space to be translated into larger distances in the 2D latent space.

### Classification performances for 19 ancestry classes

In Table 2, we report results of performances for SVM, GTM with 3 or 10 principal components, and PCA classification models based 19 ancestry classes (CEU and GBR populations were merged) from 1000 Genomes. Although the PCA plot performs rather well for the 5-classes problem, it cannot properly classify the 19 finer population classes - except for Finnish (FIN), Puerto Ricans (PUR), Peruvians (PEL), Punjabi (PJL) and Bengali (BEB). On the other hand, GTM and SVM models built from only 10 principal components can efficiently classify individuals from most of the 1000 Genomes Project populations (F1 score = 0.80). Some populations are never properly separated, even in sophisticated models taking into account more principal components; this indicates that these populations have a high genetic overlap. This is the case between the Chinese Dai (CDX) and the Kinh in Vietnam (KHV), between the Yoruba (YRI) and Esan (ESN) populations in Nigeria, and between Toscani (TSI) and Iberian populations (IBS) in Europe.

**Fig 2.**
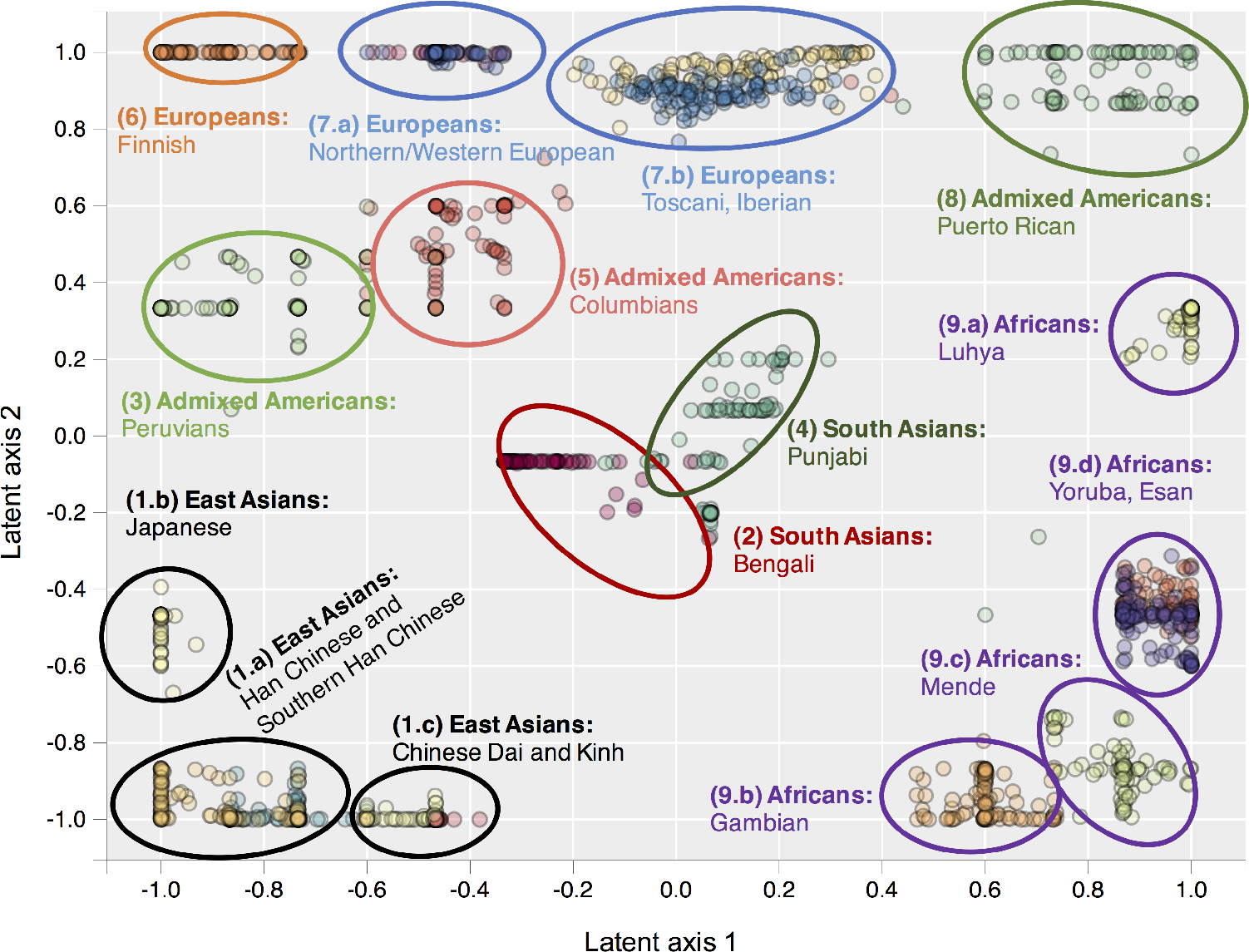
GTM with 10 principal components. Generative Topographic Mapping (GTM) plot of 20 populations from 1000 Genomes Project, built using 10 first principal components.

### Performance of classification as a function of variance explained

To investigate how the performance of 19-populations classification models (with CEU and GBR populations merged into one class) is changing depending on the percentage of variance explained, the cross-validated performance of GTM maps was evaluated by varying the number of principal components included in the model (Fig 4). The F1 score increases until it reaches a plateau around 0.80 at 10-12 principal components accounting for around 8% variance explained. Interestingly, beyond 100-200 principal components the performance starts decreasing. This could be due to including more individual-level variance, which would uncluster population structures. This indicates that the number of principal components should be optimized - our curve suggests to use between 10 and 20 components for this pruned genotype matrix.

**Fig 3.**
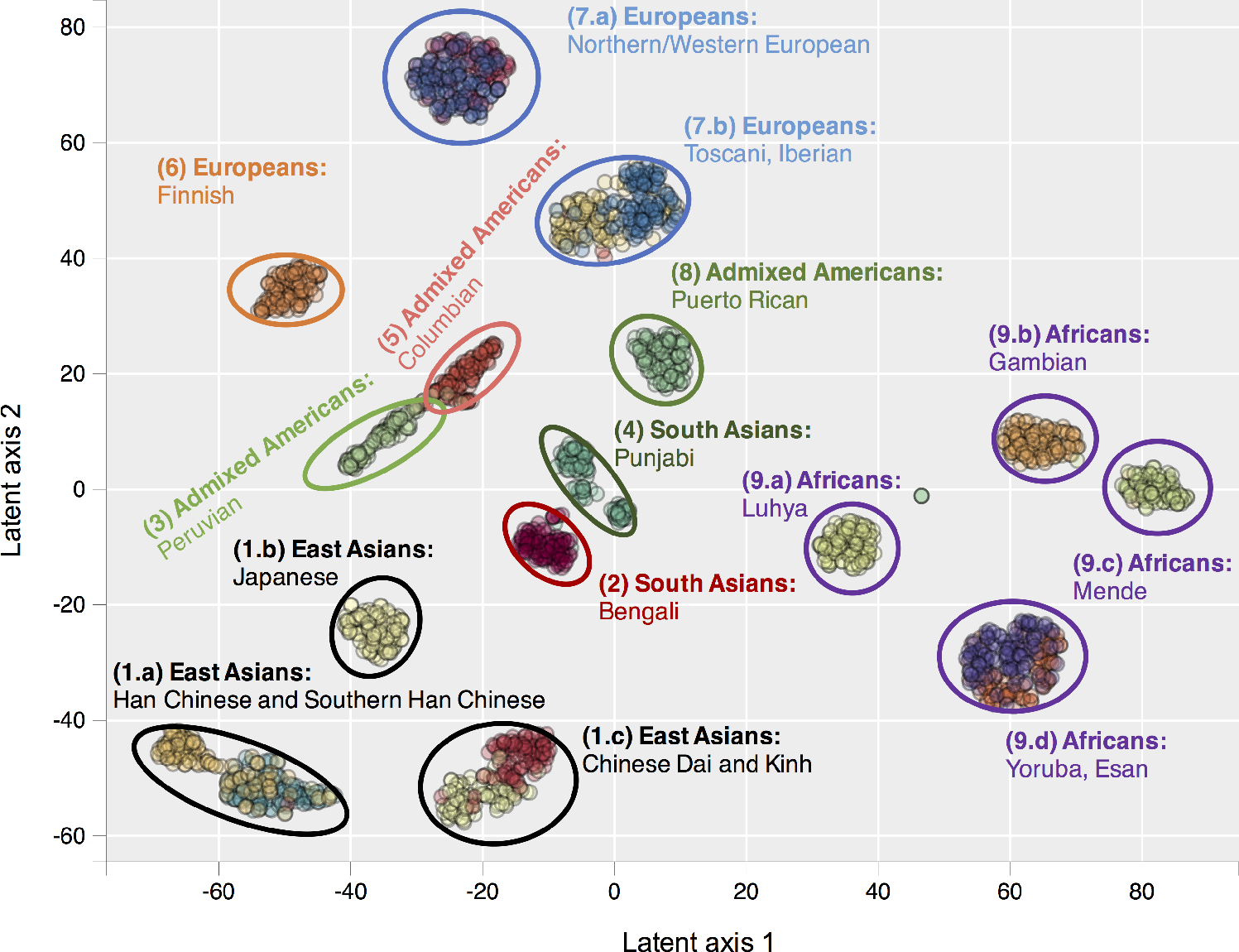
t-SNE with 10 principal components. t-distributed stochastic neighbor embedding (t-SNE) plot of 20 populations from 1000 Genomes Project, built using 10 first principal components.

**Fig 4.**
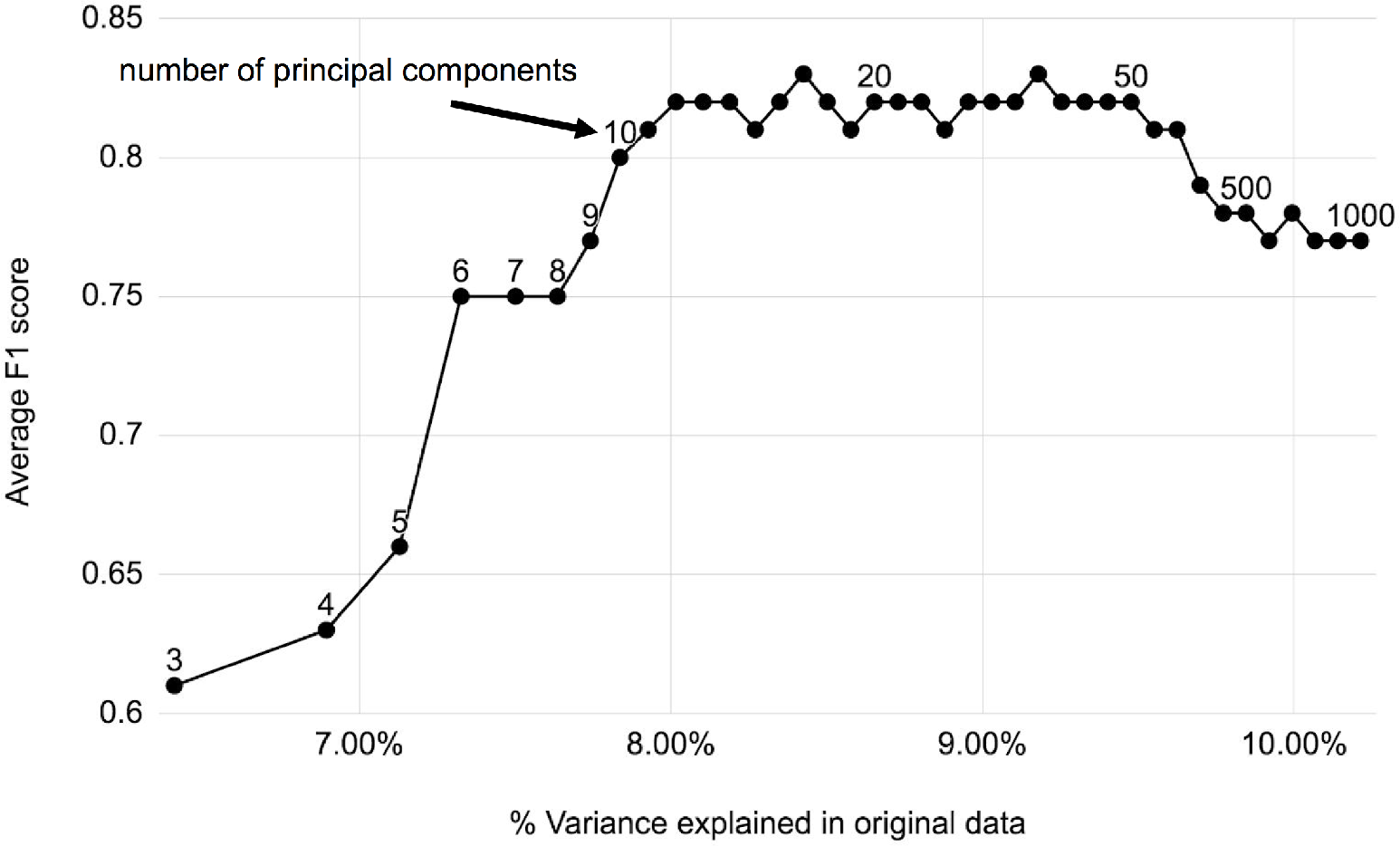
Ancestry classification performance vs. variance explained. GTM ancestry classification model performance as a function of number of principal components used to train the model.

**Table 2.** 10 times repeated 5-fold cross-validated F1 score for 19 population classes from 1000 Genomes Project using SVM, PCA or GTM.

| Ancestry | 1000G code | Population | PCA 8-NN | SVM 10 PCs | GTM 3 PCs | GTM 10 PCs |
| --- | --- | --- | --- | --- | --- | --- |
| EAS | CHB | Han Chinese | $0.20 \pm 0.01$ | $0.78 \pm 0.01$ | $0.45 \pm 0.04$ | $0.75 \pm 0.01$ |
| EAS | JPT | Japanese | $0.37 \pm 0.02$ | $1.00 \pm 0.00$ | $0.80 \pm 0.01$ | $1.00 \pm 0.00$ |
| EAS | CHS | Southern Han Chinese | $0.34 \pm 0.02$ | $0.80 \pm 0.01$ | $0.54 \pm 0.02$ | $0.80 \pm 0.01$ |
| EAS | CDX | Chinese Dai | $0.24 \pm 0.02$ | $0.10 \pm 0.02$ | $0.51 \pm 0.03$ | $0.44 \pm 0.08$ |
| EAS | KHV | Kinh in Vietnam | $0.44 \pm 0.01$ | $0.68 \pm 0.00$ | $0.63 \pm 0.01$ | $0.71 \pm 0.01$ |
| EUR | CEU+GBR | Northern/Western Eur. | $0.75 \pm 0.01$ | $0.99 \pm 0.00$ | $0.79 \pm 0.01$ | $0.99 \pm 0.00$ |
| EUR | TSI | Toscani | $0.46 \pm 0.01$ | $0.74 \pm 0.02$ | $0.58 \pm 0.01$ | $0.54 \pm 0.06$ |
| EUR | FIN | Finnish | $0.95 \pm 0.01$ | $0.99 \pm 0.00$ | $0.91 \pm 0.01$ | $0.99 \pm 0.01$ |
| EUR | IBS | Iberian | $0.35 \pm 0.03$ | $0.81 \pm 0.01$ | $0.35 \pm 0.04$ | $0.74 \pm 0.02$ |
| AFR | YRI | Yoruba in Nigeria | $0.30 \pm 0.02$ | $0.69 \pm 0.00$ | $0.15 \pm 0.03$ | $0.66 \pm 0.03$ |
| AFR | LWK | Luhya | $0.67 \pm 0.01$ | $1.00 \pm 0.00$ | $0.59 \pm 0.01$ | $1.00 \pm 0.00$ |
| AFR | GWD | Gambian | $0.26 \pm 0.02$ | $0.94 \pm 0.02$ | $0.23 \pm 0.02$ | $0.78 \pm 0.07$ |
| AFR | MSL | Mende | $0.25 \pm 0.03$ | $0.93 \pm 0.02$ | $0.35 \pm 0.03$ | $0.81 \pm 0.04$ |
| AFR | ESN | Esan in Nigeria | $0.28 \pm 0.02$ | $0.00 \pm 0.01$ | $0.19 \pm 0.05$ | $0.28 \pm 0.13$ |
| AMR | PUR | Puerto Ricans | $0.90 \pm 0.01$ | $0.86 \pm 0.02$ | $0.90 \pm 0.01$ | $0.87 \pm 0.03$ |
| AMR | CLM | Colombians | $0.69 \pm 0.01$ | $0.85 \pm 0.01$ | $0.84 \pm 0.01$ | $0.82 \pm 0.02$ |
| AMR | PEL | Peruvians | $0.88 \pm 0.01$ | $0.97 \pm 0.01$ | $0.94 \pm 0.01$ | $0.95 \pm 0.01$ |
| SAS | PJL | Punjabi | $0.89 \pm 0.01$ | $0.96 \pm 0.01$ | $0.96 \pm 0.01$ | $0.96 \pm 0.00$ |
| SAS | BEB | Bengali | $0.95 \pm 0.01$ | $0.96 \pm 0.01$ | $0.96 \pm 0.01$ | $0.96 \pm 0.01$ |
| Overall | | | $0.54 \pm 0.00$ | $0.80 \pm 0.00$ | $0.61 \pm 0.01$ | $0.80 \pm 0.01$ |

SVM 10 PCs = support vector machine classification model using 10 principal components, PCA 8-NN = k-nearest neighbours model based on 2D PCA map ( $k = 8$ ), GTM 3 or 10 PCs = bayesian classification model based on generative topographic mapping using 3 or 10 principal components. Ancestry codes: EAS = East Asians, EUR = Europeans, AFR = Africans, AMR = Admixed Americans, SAS = South Asians. CEU and GBR were merged into one class.

### Projecting populations onto the GTM map

A final map was built with 10 principal components and the complete training set of 20 populations (cf. Fig 5). The six populations that were not used to build the GTM map were used to generate posterior probabilities of superpopulation membership, which can be interpreted as the probability for a tested population *P* to belong to a superpopulation: *P*(*AFR|T*) would be the probability of African ancestry for tested population *T*. Indian Telugu from the UK (ITU), Sri Lankan Tamil from the UK (STU), and Gujarati Indian from Houston (GIH) are all predicted as South Asians (*P*(*SAS|T*) = 1) - moreover, none of them is mapped to another ancestry group. Individuals with Mexican ancestry from Los Angeles (MXL) are mostly mapped as Admixed Americans with a small European membership probability, whereas Americans of African ancestry in SW USA (ASW) and African Caribbeans in Barbados (ACB) show more mixed results - with high probabilities for both African and Admixed American superpopulations. Fig 5 shows how Americans of African ancestry in SW USA are distributed on the map: most of them are mapped near the African ancestry group but are assigned to empty nodes, where no African individual in the training set was mapped; some others are close to the Colombian/Peruvian group (AMR 1) and others to the Puerto Rican group (AMR 2).

**Fig 5.**
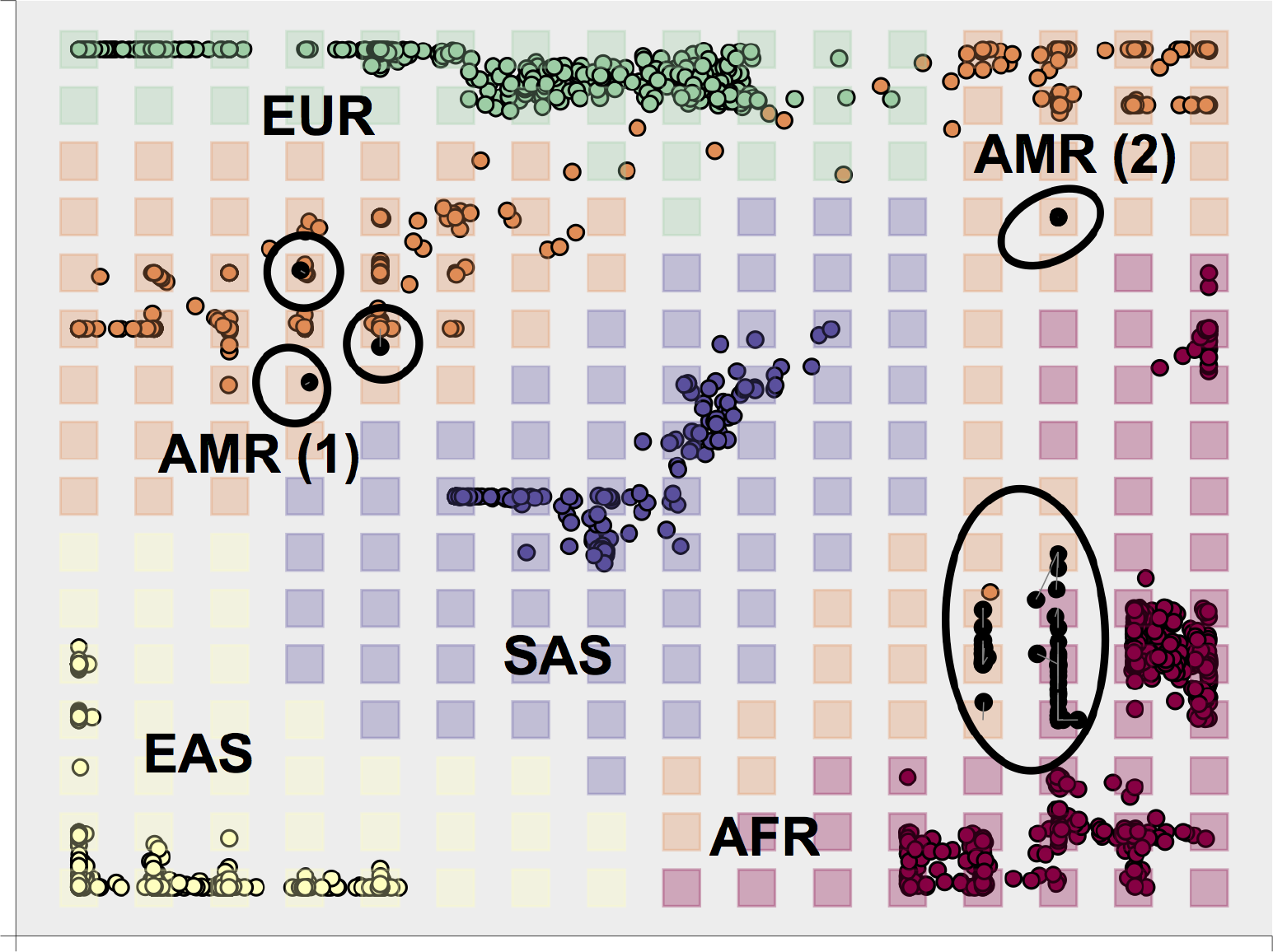
Projected Americans of African ancestry in SW USA (ASW) on a GTM map. GTM map trained with 10 principal components. Colored points represent individuals colored by ancestry or superpopulation (AFR, AMR, EAS, EUR, SAS). Squares represent GTM nodes colored by most probable ancestry. The black points in circles represent mean positions of ASW individuals projected onto the map. Grey lines map mean positions of individuals on the map to their most probable node.

**Table 3.** Posterior probabilities of superpopulation memberships in 5 test populations.

| Population (P) | $P(AFR\vert P)$ | $P(AMR\vert P)$ | $P(EAS\vert P)$ | $P(EUR\vert P)$ | $P(SAS\vert P)$ |
| --- | --- | --- | --- | --- | --- |
| Americans of African Ancestry in SW USA | 0.55 | 0.45 | 0 | 0 | 0 |
| African Caribbeans in Barbados | 0.89 | 0.11 | 0 | 0 | 0 |
| Mexican Ancestry from Los Angeles USA | 0 | 0.98 | 0 | 0.02 | 0 |
| Gujarati Indian from Houston, Texas | 0 | 0 | 0 | 0 | 1 |
| Sri Lankan Tamil from the UK | 0 | 0 | 0 | 0 | 1 |
| Indian Telugu from the UK | 0 | 0 | 0 | 0 | 1 |

Posterior probabilities of superpopulation membership obtained by GTM for 5 populations. These populations were not used to build the map; posterior probabilities using a 5-superpopulation classification model: AFR, AMR, EAS, EUR, and SAS. Ancestry codes: EAS = East Asians, EUR = Europeans, AFR = Africans, AMR = Admixed Americans, SAS = South Asians.

## Conclusion

PCA provides a good visualization of the superpopulations in the 1000 Genomes Project (AFR, AMR, EUR, EAS, SAS), but is not ideal for more fine-grained clustering and does not provide probabilistic models for admixed populations. On the other hand, both t-SNE and GTM provide a way to cluster and visualize more complex population substructures. t-SNE is arguably visually more attractive, but GTM can be harnessed to generate comprehensive ancestry classification models. Moreover, new individuals can be directly projected onto a pre-constructed GTM map - which makes it the ideal choice to cluster individuals based on pre-defined panels. We showed how to assess ancestry membership probabilities using GTM and interpret them through visualization. By generating t-SNE or GTM maps with increasing number of principal components, we can estimate the percentage of variance explained to identify population substructures - this could also be useful to account for population stratification in genome-wide association studies.

## Supporting information

**S1 Fig. GTM map of twenty 1000 Genomes Project populations.** Interactive GTM map of twenty 1000 Genomes populations. **Link to html plot here.**

**S2 Fig. t-SNE map of twenty 1000 Genomes Project populations.** Interactive t-SNE map of twenty 1000 Genomes populations. **Link to html plot here.**

**S3 Fig. GTM projection, test set 1: Americans of African ancestry in SW USA (ASW).** Projection of Americans of African ancestry in SW USA (black points) onto a GTM map trained with 10 principal components. **Link to html plot here.**

**S4 Fig. GTM projection, test set 2: African Caribbeans in Barbados (ACB).** Projection of African Caribbeans in Barbados (black points) onto a GTM map trained with 10 principal components. **Link to html plot here.**

**S5 Fig. GTM projection, test set 3: Mexican Ancestry from Los Angeles USA (MXL).** Projection of individuals of Mexican ancestry from Los Angeles USA (black points) onto a GTM map trained with 10 principal components. **Link to html plot here.**

**S6 Fig. GTM projection, test set 4: Gujarati Indian from Houston, Texas (GIH).** Projection of Gujarati Indian from Houston (black points) onto a GTM map trained with 10 principal components. **Link to html plot here.**

**S7 Fig. GTM projection, test set 5: Sri Lankan Tamil from the UK (STU).** Projection of Sri Lankan Tamil from the UK (black points) onto a GTM map trained with 10 principal components. **Link to html plot here.**

**S8 Fig. GTM projection: Indian Telugu from the UK (ITU).** Projection of Indian Telugu from the UK (black points) onto a GTM map trained with 10 principal components.

**S1 Table. 1000 Genomes Project populations.** Table of 1000 Genomes Project populations and superpopulations and the number of individuals in each category. **Link to html table here.**

**S2 Table. Variance explained in first principal components of genotype matrix.** Variance explained in 100 first principal components of the genotype matrix for twenty 1000 Genomes Projects Populations, which were used as a training set to build our models. **Link to html table here.**

**S3 Table. 5-fold cross-validated precision for twenty 1000 Genomes Project populations (19 classes) using SVM, PCA or GTM.** Precision of optimized models for the following algorithms: SVM 10 PCs = support vector machine classification model using 10 principal components, PCA 8-NN = k-nearest neighbours model based on 2D PCA map (k = 8), GTM 3 or 10 PCs = bayesian classification model based on generative topographic mapping using 3 or 10 principal components. **Link to html table here.**

**S4 Table. 5-fold cross-validated recall for twenty 1000 Genomes Project populations (19 classes) using SVM, PCA or GTM.** Recall of optimized models for the following algorithms: SVM 10 PCs = support vector machine classification model using 10 principal components, PCA 8-NN = k-nearest neighbours model based on 2D PCA map (k = 8), GTM 3 or 10 PCs = bayesian classification model based on generative topographic mapping using 3 or 10 principal components. **Link to html table here.**

**S5 Table. 5-fold cross-validated precision for five 1000 Genomes Project superpopulations (5 classes)** Precision of optimized models for the following algorithms: SVM 10 PCs = support vector machine classification model using 10 principal components, PCA 8-NN = k-nearest neighbours model based on 2D PCA map (k = 8), GTM 3 or 10 PCs = bayesian classification model based on generative topographic mapping using 3 or 10 principal components. **Link to html table here.**

**S6 Table. 5-fold cross-validate recall for five 1000 Genomes Project superpopulations (5 classes)** Recall of optimized models for the following algorithms: SVM 10 PCs = support vector machine classification model using 10 principal components, PCA 8-NN = k-nearest neighbours model based on 2D PCA map (k = 8), GTM 3 or 10 PCs = bayesian classification model based on generative topographic mapping using 3 or 10 principal components. **Link to html table here.**

## Acknowledgments

The ugtm python package used to build the GTM models was implemented by HG and accessible at https://github.com/hagax8/ugtm with a tutorial for reproducing the analyses presented here. HG and GB acknowledge funding from the US National Institute of Mental Health (PGC3: U01 MH109528). This work was also supported in part by the National Institute for Health Research (NIHR) Biomedical Research Centre at South London and Maudsley NHS Foundation Trust and King’s College London. The views expressed are those of the authors and not necessarily those of the NHS, the NIHR or the Department of Health. High performance computing facilities were funded with capital equipment grants from the GSTT Charity (STR130505) and Maudsley Charity (980).

